# DNA methylation dynamics during embryonic development and postnatal maturation of the mouse auditory organ of Corti

**DOI:** 10.1101/262832

**Authors:** Ofer Yizhar-Barnea, Cristina Valensisi, Kamal Kishore, Naresh Doni Jayavelu, Colin Andrus, Tal Koffler-Brill, Kathy Ushakov, Kobi Perl, Yael Noy, Yoni Bhonker, Mattia Pelizzola, R. David Hawkins, Karen B. Avraham

## Abstract

**Background:** The mammalian inner ear is a complex morphological structure responsible for hearing and balance, and its pathology is associated with deafness and balance disorders. To evaluate the role of epigenomic dynamics in the development and maturation of mouse inner ear sensory epithelium, we performed whole-genome bisulfite sequencing on inner ear tissue, yielding temporal base-pair resolution methylomes at key developmental time points.

**Results:** We found a late accumulation of non-CpG methylation, indicating a similarity between the inner ear sensory epithelium and neuronal tissue. Moreover, annotation of both unmethylated and low methylated regions pointed to regulatory elements active in the inner ear in proximity of and distal from transcriptional units. Finally, we identified differentially methylated regions across the transition periods. An analysis of these regions led us to identify several novel candidate regulatory factors, connecting regulatory elements from specific time points in development to molecular features that drive the development and maturation of the inner ear sensory epithelium. The GJB6 locus putative regulatory region was shown to upregulate distal GJB6 gene expression and a non-coding RNA.

**Conclusions:** Our analysis of inner ear sensory epithelium DNA methylation sheds light on novel regulatory regions in the hearing organ, and may help boost diagnostic capabilities and guide the development of therapeutics for hearing loss, by providing multiple intervention points for manipulation of the auditory system.

## Background

The mammalian inner ear is a highly complex organ that is responsible for two main functions, hearing and balance [1, 2]. Both are crucial for the survival and development of the organism throughout its lifetime. Defects in either the structure or function of any part of the inner ear sensory organs may result in auditory and vestibular impairments that represent the most diverse variety of genetic disorders [3]. The sensory organ responsible for hearing, the organ of Corti, contains the sensory epithelium (SE) [1, 2]. This region is composed of sensory cell types, i.e., the hair cells, as well as non-sensory cell types, i.e., the supporting cells. Transcriptomic approaches have been taken in order to elucidate the genetic determinants that drive each decision during the development and differentiation process of the inner ear sensory organs [4-6]. RNA high-throughput sequencing studies of the SE have led to the identification and characterization of non-coding RNAs, including microRNAs and long non-coding (lnc)RNAs [7, 8]. RNA transcriptomic analysis at single-cell level has been undertaken in this tissue [9-11], setting the stage for understanding the role of the individual cell types encompassing the SE tissue. However, to date, epigenetic analysis of the SE is still lacking, due to the technical challenges in separation of cell types and low yield of material for subsequent analysis. In recent years, technological advances in the epigenomic field have opened the door to unprecedented opportunities for unraveling of the epigenetic regulation of SE development and maturation. Transcriptomic and epigenomic analyses of the auditory system will lead to a better understanding of the genetic program of SE development and maturation and may hold the key to enable genetic manipulation and regenerative medicine in inner ear-related pathologies, including deafness.

DNA methylation remodeling is an essential component of epigenetic regulation during development [12]. This study was designed to elucidate DNA methylation dynamics during mouse SE development and maturation. To this end, we generated single nucleotide resolution genome-wide maps at key developmental time points and built a regulatory network underpinning tissue transitions throughout the developmental process, culminating in a functional, hearing inner ear. We found a close association between methylation dynamics and key developmental time points and transitions of the mammalian inner ear, with implications for the regulation of major signaling pathways such as Wnt and Notch, and mechanisms such as synaptogenesis. Finally, the analysis helped define a regulatory mechanism of *GJB6*, a critical gene in human deafness. This first reporting of a single-base resolution methylome of the mammalian inner ear has implications for regulatory pathways defining sensorineural hearing loss and deafness. Exploiting this unique resource, our study sheds new light on the pathological mechanisms of deafness in humans.

## Results

### Inner ear sensory epithelium methylomes

To explore how DNA methylation dynamics may contribute to regulate inner ear development and hearing, we generated single-base resolution methylomes of the mouse inner ear SE for three key developmental stages (Fig. 1a, Additional file 1: Figure S1a). The earliest time point investigated was embryonic day (E) 16.5, after prosensory and non-prosensory cell specification has occurred. At this time, differentiation into the organ of Corti is still ongoing, with the SE developing into a single row of sensory inner hair cells and three rows of sensory outer hair cells, surrounded by non-sensory supporting cells. The second sampled time point was immediately after birth, at postnatal day (P) 0. Although at P0 the cells are post-mitotic and the tissue can be considered non-proliferative, the organ of Corti is not yet functional and the mice cannot hear. The last time point we examined was at P22, after the mice have acquired hearing at P15 [13]. To assess DNA methylation in both the CpG (mCG) and non-CpG (mCH) context, we generated 1.3 billion unique mapped reads with an average read depth higher than 10-fold for each developmental stage (11.1X - 19.5X, Additional file 1: Figure S1b). Our genome-wide maps revealed that most of the methylated cytosines lie in the mCG context (Fig. 1b). Interestingly, at P22, we observed a prevalence of mCH methylation in both CHG and CHH, the former being 2.9% of all methylated cytosines and the latter 14.6% (Fig. 1b). While CH methylation is present in both mouse and human embryonic stem (ES) cells, it is largely lost upon cell differentiation [14, 15]. Recent evidence has shown that mCH methylation accumulates specifically in neurons during development, although its function is still unclear [16]. Thus, the increase in mCH we observed upon maturation of the SE might be associated with onset of function at both ends of the auditory system, the auditory cortex in the brain and the mechanosensory organ of Corti in the inner ear.

**Fig 1.**
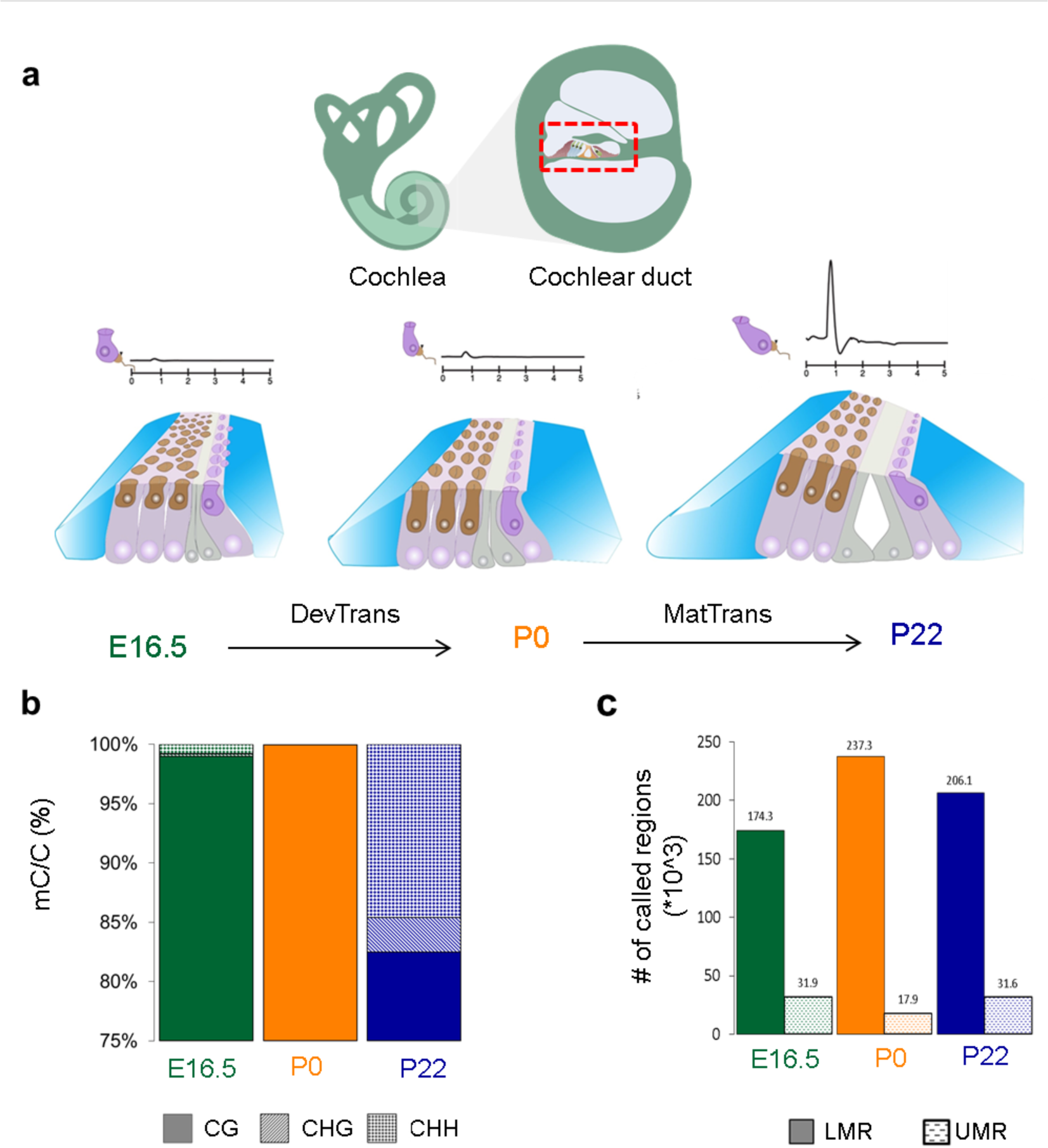
General characteristics of inner ear SE methylomes. **a** The inner ear (upper panel) is composed of the cochlea (light green) and vestibule (dark green). The former is responsible for hearing and contains the cochlear duct (red dotted line). The SE of the cochlear duct (lower panel), also known as the organ of Corti, contains sensory hair cells (brown, dark purple) and non-sensory supporting cells (blue, grey, light purple) at E16.5, P0 and P22. At the first two stages, there is no functional hearing, as demonstrated by peaks of the auditory brainstem response (ABR), while hearing is functional at P22. **b** Composition of all methylated cytosine loci in the genome of all three time points, CG dinucleotide (solid), CHG (dashed line) and CHH (dotted). **c** Number of called LMRs (solid) and UMRs (dotted) at each time point.

### Regulatory landscape of the inner ear SE

To generate regulatory maps of inner ear SE development and maturation, we identified unmethylated and low-methylated regions (UMR/LMRs), which are known to be characteristic of regulatory elements, such as promoters and enhancers [17, 18]. UMRs are defined as CG-rich regions with an average methylation lower than 10%, while LMRs are defined as regions where the average methylation ranges from 10% to 50%. UMRs are predominantly found at promoters, while LMRs are predominantly located distal to promoters in intergenic or intronic regions. Typically, LMRs tend to be enriched for H3K4me1, DNase I hypersensitive sites (DHS) and p300/CBP. We identified 31,916, 17,945 and 31,564 UMRs and 174,347, 237,282 and 206,144 LMRs in E16.5, P0 and P22 methylomes, respectively (Fig. 1c; Additional file 2: Table S1), which have the expected percent methylation and genome localization characteristics (Additional file 1: Figure S1c-h).

UMRs and LMRs provided a means to identify novel regulatory elements within the inner ear SE (Fig. 2a). UMRs were validated by their localization to annotated transcription start sites (TSS) and CpG islands (CGIs) (Additional file 1: Figure S1e, g). To ensure that LMRs were representative of known distal regulatory elements, we determined if they overlapped known DHS in the mouse genome, which are indicative of transcription factor (TF) binding. Using data from the mouse ENCODE project [19], we found that 93.6%, 85.5% and 93.3% of LMRs from E16.5, P0, and P22, respectively, where hypersensitive in other mouse tissues, confirming their status as transcription factor binding sites (TFBS) (Fig. 2b). Of LMRs overlapping known DHS, 9.8-15.5% contained a CTCF motif and overlapped known CTCF binding sites [19, 20], suggesting a role as insulators or in three-dimensional (3D) genome architecture (Fig. 2b). The remaining 84.5-90.2% overlapped known H3K4me1 sites from the mouse ENCODE project (Fig. 2b) [19, 20], suggesting they may function as enhancer elements [21].

**Fig 2.**
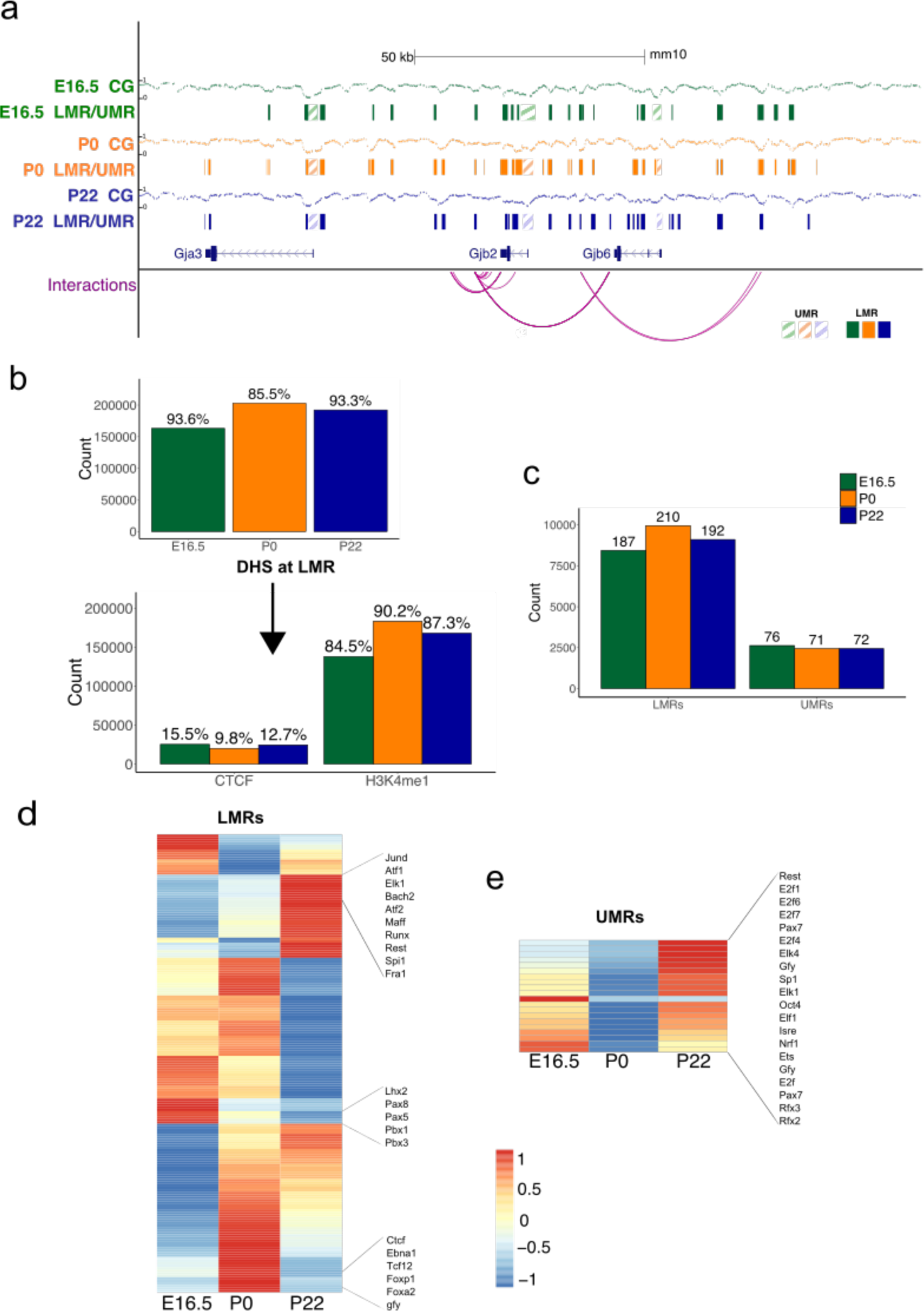
Putative regulatory landscape of the inner ear SE. **a** Browser shot at the *Gjb2* gene surrounding locus, illustrating percent methylation levels, LMRs, UMRs and their putative interactions with target genes, **b** Barplot showing the number of identified LMRs overlapping with known DHS sites. The numbers on each bar denote percent overlaps. Bottom barplot shows the number of hypersensitive LMRs overlapping with CTCF binding sites and H3K4mel enhancer peaks. The numbers on each bar denote percent overlaps, **c** Barplot showing the number of putative target genes interacting with LMRs and UMRs. The numbers on each bar denote the count of known deafness target genes, **d** Heatmap of row normalized - log(P-value) for the enriched TF motifs present at LMRs and their relative enrichment across three time points, **e** Heatmap of row normalized -log(P-value) for the enriched TF motifs present at UMRs and their relative enrichment across three time points.

To understand the regulatory implications of these elements, we determined their target genes (Fig. 2a, c; Additional file 3: Table S2). Any distal UMRs, such as those at non-promoter CGIs, and LMRs were linked to putative target genes using both known 3D interactions from C-based data (3C, 4C, 5C, ChIA-PET, or HiC) and predicted interactions based on a random forest classifier trained on known interactions and non-interactions from the 4D genome database (Fig. 2a) [22]. Importantly, we were able to link UMRs and LMRs with known mouse inner ear development, functionally relevant and known hearing impairment or deafness genes [23]. On average, 73 and 196 genes could be linked to UMRs and LMRs, respectively (Fig. 2c; Additional file 4: Table S3). Next, we determined enriched TFBS motifs within UMRs and LMRs to predict putative regulators of target genes (Fig. 2d-e; Additional file 5: Table S4). Motifs for several known inner ear TFs were found enriched within all LMRs analyzed, such as *Atoh1, Stat3, Gata3, Sox10* and *Sox2* and were also correlated with available inner ear SE transcriptomes [6, 8]. It is clear from the TFBS enrichment heatmap across time points that some motifs are enriched at one time point and not others, likely reflecting a temporal dimension to the relevance of the binding of TF and their asserted regulation. One such interesting example is the *Oct4* (also known as *Pou5f1*) TFBS, one of the key factors for proliferation and pluripotency [24], which exhibit a significantly higher enrichment at P0 LMRs compared with E16.5 and P22 (Fig. 2d). Oct4 is known to interact with other TFs mentioned previously, such as *Sox2* [25]. Another interesting TFBS motif found in LMRs is of *NF-**κ**B* (p = 1 x 10^−73^), which was previously shown to play a role in the response of the cochlea to both noise-induced damage and inflammation [26, 27]. At UMRs, we found significant enrichment for *Rfx2* and *Rfx3* enriched, both members of the RFX transcription factor family and associated with hair cells [5, 9]. We have also observed a significant enrichment of a *FoxO1* binding motif (CTGTTTAC) in LMRs across all time points, while the same motif was not found enriched at UMR locations. *FoxO1* is predominantly expressed in supporting cells of the SE and is a direct target of *miR-96*, a key non-coding regulator of inner ear SE development [28]. Overexpression of *miR-182*, part of the *miR* triad of *−96, −182* and *-183*, inhibits *FoxO1* translation [29] and ER-stress induced damage to the inner ear SE [30]. Interestingly, we found that a *Bach2* motif was significantly enriched across all time points (E16.5, P0 and P22, p = 1x10^−9^, 1x10^−16^, 1x10^−76^, respectively). This TF plays a role in the Bcl6-Bcl2-p53 axis in controlling apoptosis in several non-inner ear tissue types (reviewed in [31]), as well as has late onset expression in the developing chick ridge, otic cup and/or neural crest [32]. However, *Bach2* has not been reported to play a role in the inner ear development.

### Differential methylation during development and maturation transitions

To explore how DNA methylation dynamics may contribute to regulate inner ear SE development and onset of hearing, we focused on two sequential transitions. The first is the development of the inner ear SE, taking place during the transition between E16.5 to P0; thus, we refer to this transition as the “developmental transition" (DevTrans). The second is the maturation of inner ear SE, from P0 to P22. Given that hearing is initiated at postnatal day P12, and is functional by P15 [13], we refer to this period as “maturation transition" (MatTrans) (Fig. 1a). For each transition, we called differentially methylated regions (DMRs), whereby at least a 30% change in methylation (p-value < 0.05) occurred between the start and end time point, resulting in loss of methylation (hypo-DMR) or gain in methylation (hyper-DMR) during the transition (Fig. 3a, Additional file 6: Fig. S2a, Additional file 7: Table S5). While the number of hypo- and hyper-DMRs were similar for each transition period (DevTrans: 998 vs 1340; MatTrans: 4470 vs 5540), we observed that the DNA methylation signatures change more during the MatTrans period than during DevTrans. DMRs appeared to have a similar size, between 661 to 773 bp, across both transitions (Additional file 6: Fig. S2b) and to be largely located in intronic and intergenic regions (Additional file 6: Fig. S2c). Consistent with their genomic location, DMRs were more frequently found to overlap LMRs than UMRs (Additional file 6: Fig. S2d).

**Fig 3.**
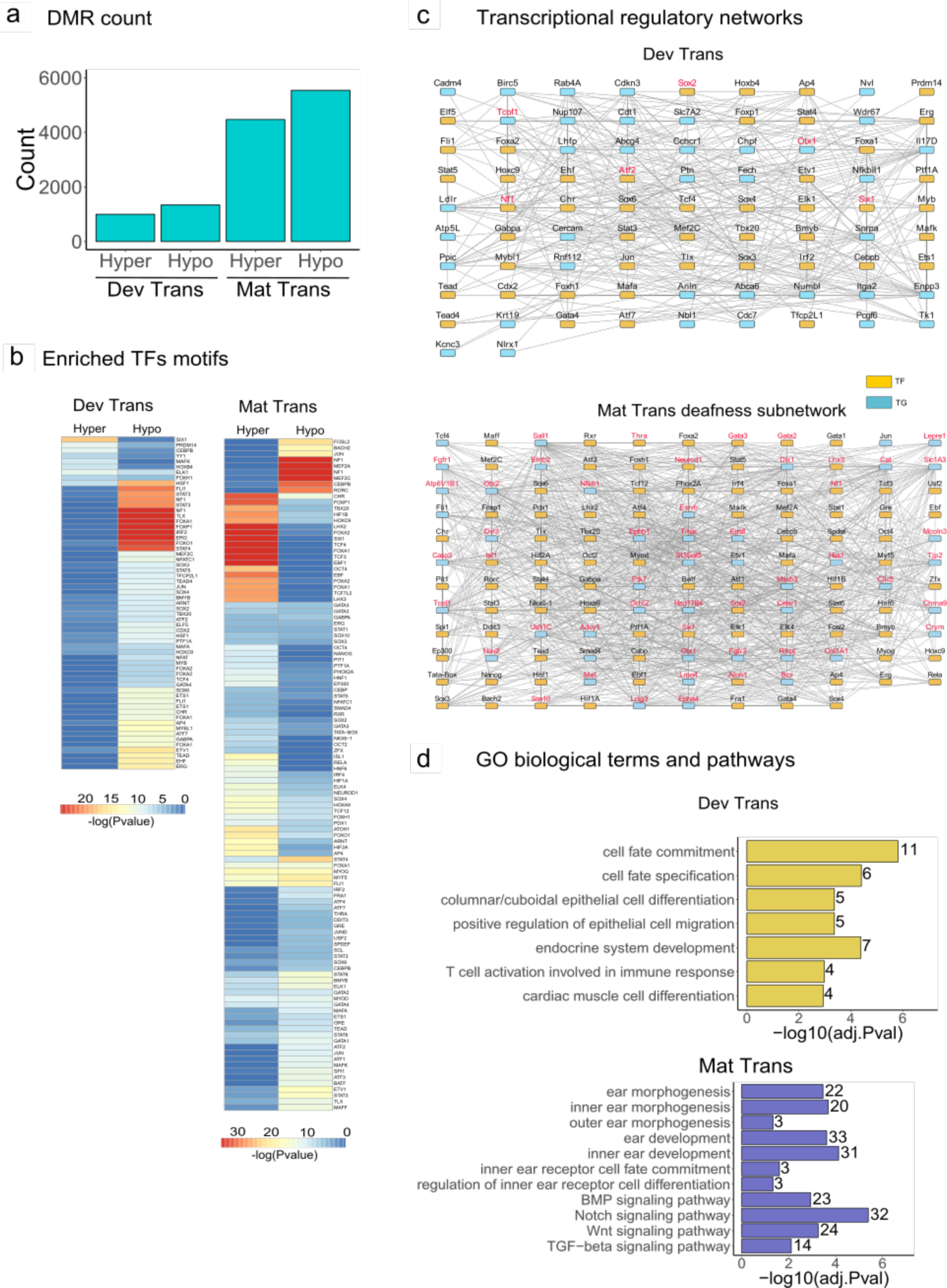
Identification of differentially methylated regions during development and maturation. **a** Barplot showing number of identified hyper- and hypo-DMRs during DevTrans and MatTrans. **b** The enriched TF motifs present at DMRs. **c** Top: *In silico* transcriptional regulatory network based on DMR target gene interactions during DevTrans. Known deafness TFs/genes are shown in red colored text. Bottom: The subnetwork based DMR target gene interactions of known deafness genes during MatTrans. The deafness associated TFs/genes are shown in red colored text. See Additional file 8: Figure S3 for complete MatTrans network. **d** The enriched GO biological terms and pathways for DevTrans (top) and MatTrans (bottom).

We subjected all DMRs to TFBS motif analysis to obtain a better understanding of how differential methylation might impact gene regulation (Fig. 3b, Additional file 8: Fig. S3a; Additional file 5: Table S4). A number of known inner ear regulators were enriched, including *Six1* [33], *Stat3* [34], *Pou3f4* [35, 36] and Wnt factors *Tcf3* and *Tcf4* [37-39]. *FoxO1*, which was enriched in LMRs, was found to be enriched at DevTrans hypomethylated DMRs and both hypo- and hypermethylated DMRs at the later transition. Among other factors found in both LMRs and DMRs is *Sox2*, a key factor in the prosensory differentiation of hair cells [40]. Interestingly, the motif for *Atoh1* becomes methylated by P22 (MatTrans hyper-DMRs), coinciding with down-regulation of *Atoh1* after P5, and it early role in major cell type differentiation processes [41]. The *Bach2* motif (TGCTGAGTCA) was enriched only at MatTrans hypomethylated DMRs (p = 1*10^−5^) to coincide with the highest enrichment score exhibited in P22 LMRs for this TFBS.

Most DMRs are distal to genes they likely regulate; as a result, we determined DMR putative target genes during both transitions as we did for LMRs. We leveraged the DMR motif analysis of TFs and putative target genes (TGs) to construct DevTrans and MatTrans regulatory networks (Fig. 3c, Additional file 8: Fig. S3; Additional file 9: Table S6). These networks illustrate how regulatory elements that exhibit changes in DNA methylation might alter TF binding to control gene expression of their target genes. Each network also implicates known mouse deafness genes for additional insight on how these genes are regulated. Many known deafness genes are TFs. The network analysis is predictive of how a cascade of genes become altered when these TFs are mutated in mice. We conducted Gene Ontology (GO) term enrichment analysis for each network based on TFs and TGs (Fig. 3d; Additional file 10: Table S7). The DevTrans network was enriched for terms such as ‘cell fate commitment’, ‘cell fate specification’, and ‘epithelial cell differentiation’. The MatTrans network, which included substantially more DMRs and therefore TGs, included various terms that included ‘ear morphogenesis’ and ‘ear development’. This network was also enriched for BMP, Notch, Wnt, and TGF-beta signaling pathways.

### Time point-specific DNA methylation changes

Changes in DNA methylation are subject to thresholds that can be arbitrary. When we compared our UMR/LMR calls across time points to DMR calls, which required a 30% methylation change, our results indicate that a larger portion of LMRs, and to a certain extent UMRs, appear to be more dynamic than our DMR analysis revealed (Additional file 11: Figure S4A). Because small changes in the average methylation of UMRs and LMRs could alter a specific TFBS, and because the percent methylation calls are on bulk tissue, we sought to expand our analysis. We determined 5692 UMRs and 314,192 LMRs identified in a single time point as additional sites of interest. These are referred to as time point-specific UMRs and LMRs (Fig. 4a).

**Fig 4.**
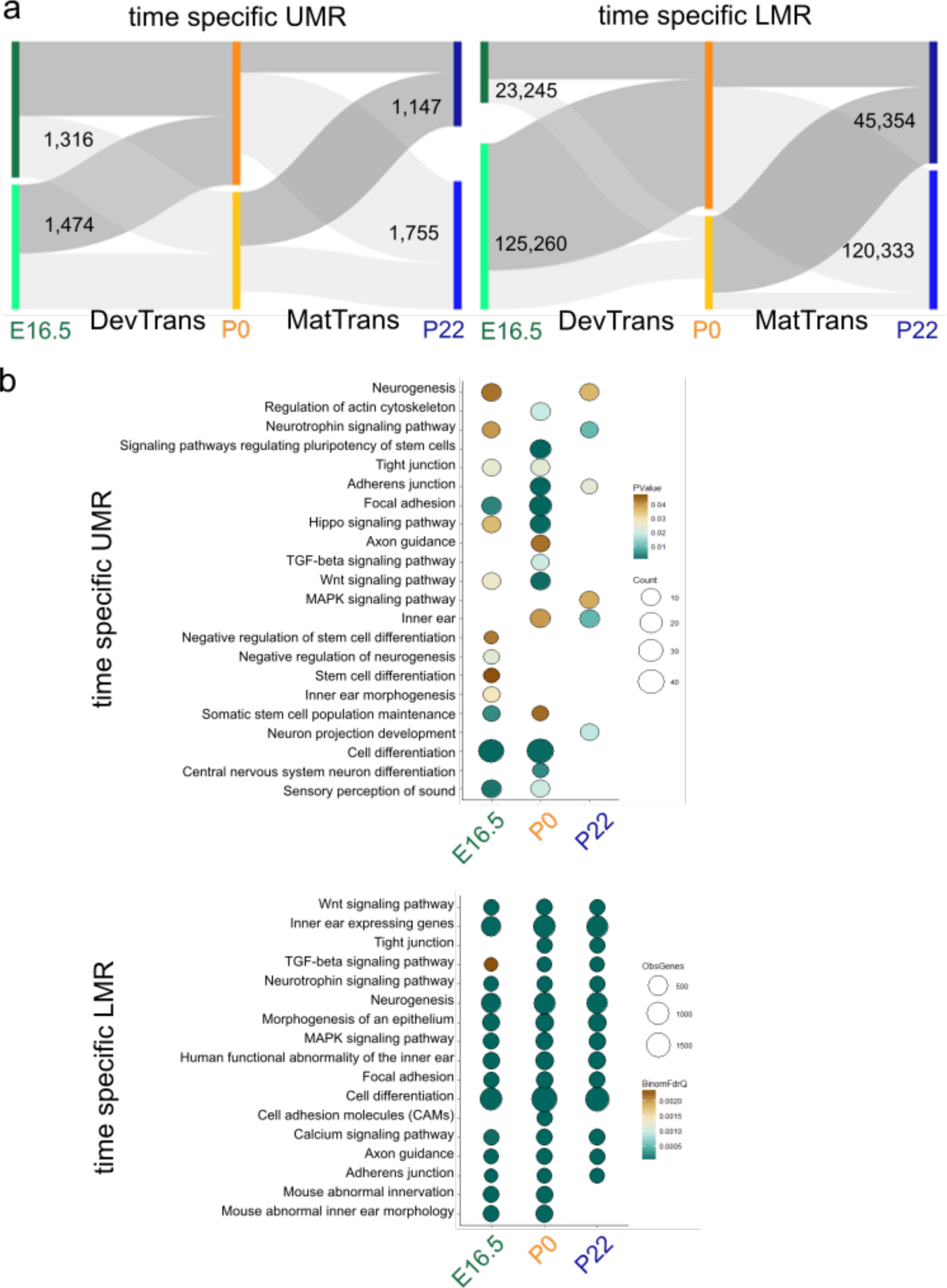
Analysis of UMR and LMR putative regulatory regions. **a** Time point-specific UMRs (left) and LMRs (right) and their dynamics through the time points (numbers indicating unique region), represented as sankey plots; regions defined at a specific time point as UMR/LMR (dark grey) and regions not defined as UMR/LMR (light grey). **b** GO term enrichment analysis of time point-specific UMRs (top) and time point-specific LMR (bottom) associated genes; for UMR analysis, circle fill is according to p value (<0.05) and for time point-specific LMR according to q value (FDR < 0.05). Circle diameter is proportional to the number of genes associated to terms.

We then performed GO term enrichment analysis for the genes associated with time point-specific UMRs (Fig. 4a-b; Additional file 11: Figure S4a; Additional file 12: Table S8). The results revealed a number of tissue-specific terms that are enriched, such as ‘sensory perception of sound’ at E16.5 and P0, ‘inner ear morphogenesis’ at E16.5, and ‘inner ear’ hypo-methylated at P0 and P22. We also observed signaling pathways known to govern inner ear SE development, such as Wnt, neurotrophins or TGF-beta signaling pathways^1,2^. In addition, many neuronal tissue-related terms (e.g. ‘neurogenesis’, 'axon guidance') were found enriched at all time points. As the SE goes through a developmental transition from E16.5 to P0, we were not surprised to find ‘cell differentiation’ enriched at both earlier time points, but lacking in the more mature SE at P22. We observed ‘cell differentiation’ and ‘morphogenesis of epithelium’ to be enriched (Fig. 4b). ‘Inner ear’ expressed genes were enriched within the putative target genes of the time point-specific LMR at all time points, as well as the human phenotype of ‘functional abnormality of the inner ear’, the ‘abnormal inner ear morphology’ and ‘abnormal innervation’ phenotypes. As with time point-specific UMRs, the putative target genes of the time point-specific LMR GO analysis were enriched for various signaling pathways such as Wnt, neurotrophins and calcium signaling, and ‘neurogenesis’ was one of the leading biological processes to be enriched.

Time point-specific LMRs and UMRs were enriched for immune-related response genes. Time point-specific LMRs were enriched for broad 'immune system development' terms and mouse immune phenotypes (Additional file 12: Table S8; Additional file 13: Table S9), while time point-specific UMRs were enriched for immune-specific pathways, such as the ‘TGF-beta signaling pathway’ (Fig. 4b; Additional file 12: Table S8; Additional file 13: Table S9).

### Gene expression and DNA methylation

We compared available transcriptomic data available for SE [6, 8] to the DMR dynamics at genes promoters (Fig. 5a-b) and genebodies (Fig. 5c-d). We did not observe a negative correlation trend between the expression of genes with methylation status dynamics at their promoter and their differential expression across transitions. This was also true when we focused on genes with CpG islands overlapping their promoter (Fig. 5c) or used the available Shared Harvard Inner-Ear Laboratory Database (SHIELD) [6] (Additional file 14: Fig S5a-d). In order to relate to the interaction of DNA methylation and the expression level of genes, we focused on genes that exhibit an expected correlation between gene expression in our RNA seq data [8] and the DevTrans DMRs (Fig. 5d); specifically, the negative correlation between DNA methylation of CpG on gene promoter and the gene’s differential expression and positive correlation between genebody DNA methylation and gene expression [42-44]. Out of 2338 DevTrans DMRs, only 334 DMRs were residing in gene’ promoters, and 1259 were residing in genebodies. Two hundred and eleven genes presented anti-correlation between promoter DNA methylation and gene expression (Fig. 5d) and 180 genes presented with a positive correlation between DNA methylation at genebodies’ and the gene expression. Among the genes listed, we found *Otx1*, a transcription factor required for inner ear development. *Otx1* expression is downregulated (-1.38-fold decrease, FDR= 1.98*10^−9^), while it promoter is hypermethylated during DevTrans (%MethDiff = 24.195%, p = 0). *Otx1* expression level had been considered in the literature to be low to none at E14-E15 in the mouse [45]. Another interesting gene, *Espn*, was shown to go through dual regulation over its expression (1.44-fold increase, FDR=1.43*10^−11^), through its DNA methylation dynamics of both promoter (%MethDiff = −48.45%, p = 0) and genebody (%MethDiff = 34.04%, p = 0) DNA methylation.

**Fig 5.**
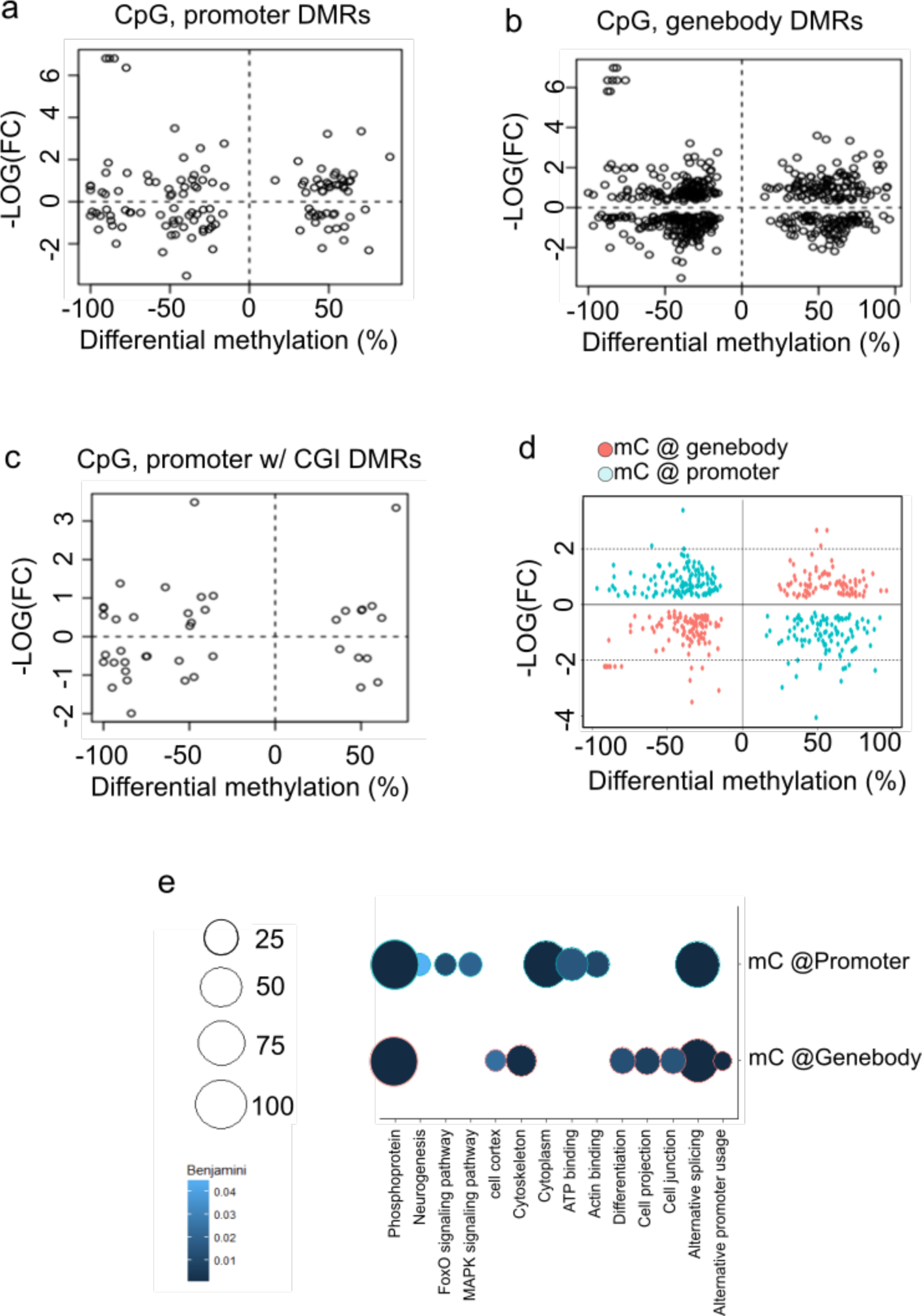
Analyzing the correlation between DNA methylation dynamics and transcriptome of the inner ear SE. Scatter plot of DevTrans SE RNA-seq expression data [8] plotted against the gene associated DMR differential methylation score. **a** DMRs analyzed in the CpG context and residing in gene promoters. **b** DMRs in gene promoter that overlap CpG islands. **C** DMRs in the CpG context found in the body of the gene. **d** Focusing on genes presenting positive correlation of gene expression and genebody methylation dynamic (turquoise), and genes presenting negative correlation between gene expression and genes’ promoter DNA methylation dynamic (red). **e** GO term enrichment analysis of either gene subsets, indicated as ‘genebody’ or ‘promoter’; circle fill is according to Benjamini score (<0.05) and Circle diameter is proportional to the number of genes associated to terms.

We performed GO analysis on the 211 and 180 genes described above (Fig. 5e; Additional file 15: Table S10), and observed molecular function such as ‘alternative splicing’ to come up in both gene lists (Benjamini = 2.56*10”^14^), as well as ‘actin binding’ (Benjamini = 0.01) within promoter methylated genes. Interestingly, the 211 genes presenting with anticorrelation between promoter methylation and gene expression dynamics were found enriched for the KEGG pathway ‘FoxO signaling pathway’ (Benjamini = 0.012), linking back to the *FoxO1* TFBS described previously.

### Mouse inner ear SE LMRs are informative of regulation of human deafness

The annotation of *cis*-regulatory elements in the human genome has increased our understanding of disease-associated variants by revealing their location in regulatory elements such as enhancers [46-49]. We investigated how mouse inner ear SE LMRs could expand our understanding of human deafness by annotating putative regulatory regions. We converted our mouse (mm10) LMR coordinates to human (hg19) coordinates using UCSC liftover (E16.5: 56,439; P0: 83,299; P22: 67,545 LMRs lifted over). On average, 84% of mouse-to-human LMRs are both hypersensitive (DHS) and marked by H3K4me1 in at least one human cell or tissue type, supporting the assumption that these elements act as enhancers in the human genome. Next, we determined if hearing-related variants from GWAS and their proxy SNPs (https://www.ebi.ac.uk/gwas/) in linkage disequilibrium (LD) overlapped our mouse-to-human LMRs. Although most hearing-related studies have limited associated variants, we found 24 variants associated with age-related hearing impairment, 21 with ear morphology, and two with hearing impairment at LMRs. All but four of the LMRs are marked by H3K4me1 in at least one human cell or tissue type, suggesting they are enhancer elements. We were able to assign 12 variants to target genes using known and predicted 3D interactions in human cells, as described for mouse. Many of these interacting variants are clustered on chromosome 6, primarily in 6p21 and 6p22 (Fig. 6a). Chromosome 6 is a hotspot of non-syndromic deafness genes, harboring seven autosomal recessive genes or loci (four on 6p21), and six autosomal dominant genes or loci (three on 6p21). Two autosomal dominant loci, DFNA21 (6p21) and DFNA31 (6p21.3) remain undetermined [50, 51]. We also examined the regions associated with deafness loci with unknown genes described in the Hereditary Hearing Loss Homepage (http://hereditaryhearingloss.org/). Chromosomal regions, derived from microsatellite markers or coordinates of cytogenetic bands, were located in the relevant manuscripts, identified in hg19, and converted to the homologous mouse interval (mm10) [8]. We identified SE LMRs in all of these regions, suggesting they may be associated with deafness in humans. These mouse-to-human maps provide a valuable resource for annotating putative regulatory elements relevant to the genetics of human deafness.

**Fig 6.**
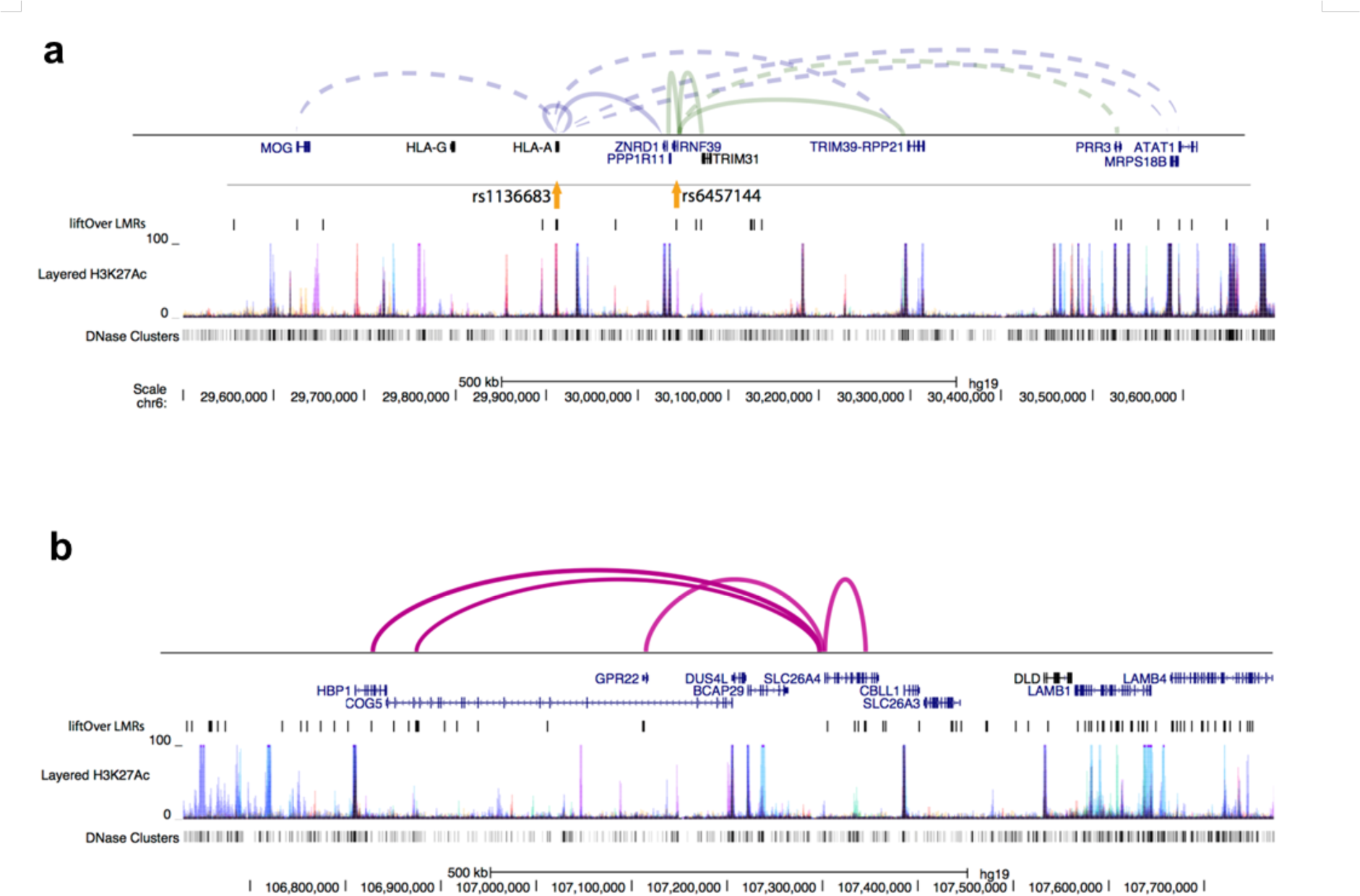
Inner ear SE LMRs are informative about human deafness. **a** Browser shot of hearing-related variants at mouse-to-human LMRs and their target gene interactions on chromosome 6p21. Known interactions are shown as solid lines, predicted interactions as dashed lines. **b** Browser shot of mouse-to-human LMRs and known interactions around the Pendred syndrome gene *SLC26A4.* For **a** and **b**, layered H3K27ac from seven cell types and DHS data are shown to illustrate how SE LMRs can guide identification of c/s-regulatory elements of interest.

A number of affected individuals with hereditary hearing impairment do not harbor known causative, coding variants. For example, almost half the cases of Pendred syndrome, a disorder that includes hearing impairment due to inner ear malformations in the cochlea and enlarged vestibular aqueduct (EVA), are caused by mutations in *SLC26A4* [52]. For the remaining half, the cause is largely unknown. It is feasible that some manifestations of hearing impairment are caused by distal regulatory variants. Mapping mouse LMRs from the inner ear SE to human coordinates could distinguish which of the known human *cis*-regulatory elements to focus on for regulation of known deafness genes. We have illustrated such an example for *SLC26A4*, where several mouse-to-human LMRs are near the gene and at least four have known interactions (Fig. 6b).

### The human GJB2-GJB6 locus harbors a putative enhancer modulating GJB6 expression

In order to validate mouse-to-human LMRs predicted to operate as gene enhancers, we utilized the CRISPRon system, based on a mutated (“dead”) Cas9 protein (dCas9) that is unable to exert a DNA double strand cut or single strand nicking, to which the 10x repeats of the herpes simplex virus activation domain (VP16) is attached (dCas9-VP160) [53]. As our top candidate for validation, we chose a *GJB2* proximal regulatory region 1.2kbp upstream of the *GJB2* TSS (Fig. 7a), which is in the region between *GJB2* and *GJB6.* Both are two prominent deafness genes, and *GJB2* is associated with approximately 30% of deafness in the Israeli Jewish population [54, 55] and up to 50% of deafness in other populations [56]. According to ENCODE data, our candidate region is clearly marked as an ‘active enhancer’ by ChromHMM segmentation in a *GJB2* and *GJB6* expressing cell line, NHEK (Normal Human Epidermal Keratinocytes) (Fig. 7a), confirming our *in silico* prediction of this region as a putative enhancer in the inner ear SE. Two guide RNAs (gRNA) were designed to direct the dCas9- VP160 to the *GJB2* proximal enhancer (Fig. 7a), and transfected into HEK293T cells that neither express *GJB2* or *GJB6.* Using either guide, or their combination, resulted in no modulation of *GJB2* expression (Fig. 7b), despite the proximity to the *GJB2* promoter region. Surprisingly, using gRNA#1 alone resulted in a 3.36-fold increase in *GJB6* expression levels (p = 0.0043, n=4), while gRNA#2 had no significant (p >0.05) effect and the combination of both significantly increased *GJB6* expression levels, but to a lower extent than gRNA#1 alone (1.37-fold, p=0.035, n=4). As a negative control, we used gRNA targeting TetO. Interestingly, we observed that the modulation of *ATOH1*, performed as a positive control for the CRIPSRon system, actually enhanced *GJB6* expression and had no effect on *GJB2* expression (Fig. 7b). Also, targeting the putative enhancer with the potent gRNA#1 increased *ATOH1* expression by 1.27-fold (p= 0.047, n=4). We tested the expression of a non-coding RNA, *RP11-264J4.10*, found in close proximity to the candidate regulatory region (Fig. 7a), and observed the same pattern as for *GJB6* expression modulation, namely gRNA#1 increased the ncRNA expression by 5.87-fold (p = 0.024, n=5) and *ATOH1* expression activation increased it by 10.3-fold (p = 0.042, n=5) (Fig. 7c), suggesting the enhancer interacts with multiple genes. Analyzing the candidate regulatory region (+/-60bp on each side) for ATOH1 binding motifs using JASPAR (http://jaspar2016.genereg.net) [57] resulted in one predicted binding site (GACAGATTTG, Score = 4.257, Fig. 7d), which may explain how *ATOH1* activation increases *GJB6* and *RP11- 264J4.10* expression. Our data suggests that by lifting over mouse LMR putative enhancers to the human genome, we have annotated a human relevant enhancer region, proximal to the *GJB2* gene and regulating the distal *GJB6* gene and a previously annotated ncRNA.

**Fig 7.**
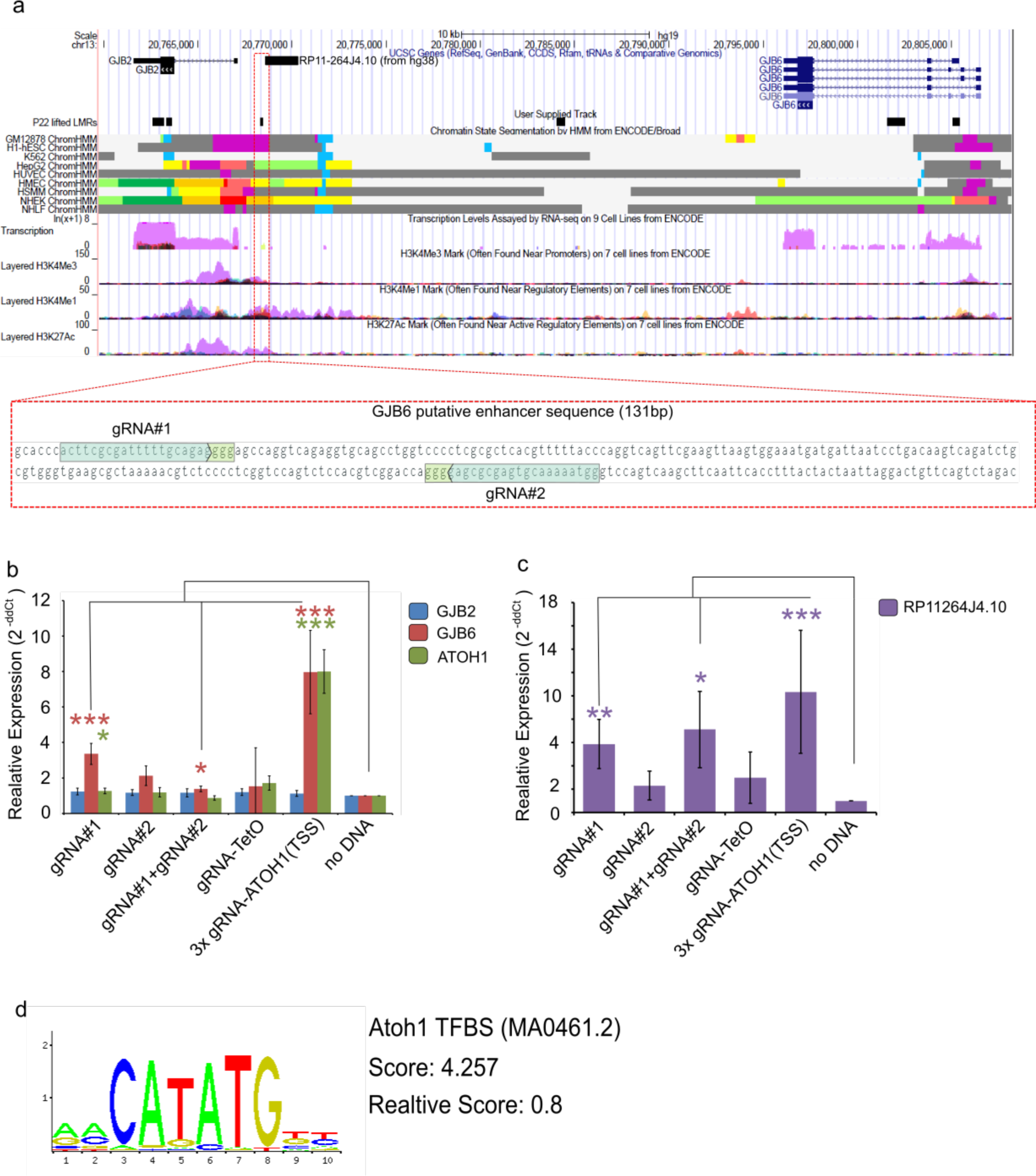
CRISPRon based modulation of GJB6 and RP11-264J4.10 (ncRNA) expression via a putative enhancer. **a** UCSC browser snapshot presenting the candidate enhancer lifted over from mouse (mm10) to human (hg19) (track: “P22 lifted LMRs”, black box indiactors); additional information derived from ENCODE regulation hub is presented to support the characterization of the candidate sequence to act as enhancer (track:‘transcription’ - purple signal = NHEK cell line, track:‘NHEK ChromHMM’ - orange indicating ‘Active enhancer’). Below, zoom in on the candidate sequence and the gRNA layout within it. Expression of *GJB2, GJB6* and *ATOH1* (b) and *RP11-264J4.10* ncRNA (c) measured qRT-PCR (normalized to *GAPDH*, compared to ‘no DNA’, n=4-5, *** p <0.01, * p<0.05). **d** ATOH1 transcription factor binding motif found on the edges of the candidate enhancer, score is calculated by JASPAR.

## Discussion

Given the critical role that DNA methylation is known to play in development and disease, we took a whole organ view of DNA methylation dynamics and the annotation of regulatory elements according to DNA methylation status, by performing a whole-genome analysis of the inner ear cochlear SE at single-base-pair resolution. While the mouse methylome has been studied in detail [16, 18, 58], inner ear tissues have not previously been examined at this resolution. The cochlear SE is the sole sensory organ responsible for the translation of air vibrations into auditory signals that are then relayed to the brain to allow mammals to hear. Our approach to analyze the inner ear SE as a whole organ reflects the DNA methylation status of all cellular populations comprising this complex tissue, restricting our conclusion to the organ of Corti as a single, complete unit. We have demonstrated, for the first time, the genome-wide involvement of DNA methylation and its dynamics in the mouse inner ear SE development from E16.5, following exit of the cell cycle but prior to full differentiation of hair and supporting cells, to maturation at P22, representing the fully functional tissue after hearing onset. Variations during these transitions include differences in the CG (mCG) and non-CpG (mCH) methylation status, dynamic usage of putative regulatory regions through identification of UMRs and LMRs, as well as regions of differential methylation during development and maturation. We also tested the relationships between DNA methylation and gene expression in this highly complex tissue. The analysis enables us to highlight a number of genes, and biological and molecular functions, some of which have been previously characterized or previously annotated in the inner ear but with unknown function, and to define regulatory regions characterized by methylation status that are important for the development and maturation of the SE.

### Inner ear sensory epithelium methylomes are unique in comparison to other non-neuronal tissue

Non-CpG methylation, in the context of CHG and CHH, characterizes pluripotent ESC and distinct fully differentiated brain cell types [14, 16, 59]. However, the prevalence of non-CpG methylation in other somatic tissue types is very low [15, 60]. The inner ear SE methylome revealed a sharp increase in non-CpG methylation at the later developmental time point, P22. The proportions of both mCHG and mCHH increased when compared with either of the earlier time points, reaching as high as 14% of mC in the context of CHH. Non-CpG methylation plays a key role in establishing neuroplasticity in the adult differentiated brain [59]. CHG/CHH methylation is accumulated in certain types of brain neurons as part of neuronal circuit maturation [16]. This process of maturation begins after P7, and reaches a peak in 4-week-old mice, parallel to inhibitory synaptogenesis in the mouse somatosensory cortex [61]. Interestingly, mice only begin to hear at P12, with a fully functional hearing organ by P15 [13]. Earlier in the development of the inner ear, parcellation of the early otic vesicle into a non-sensory, neurogenic organ and either to an auditory or vestibular mechanosensory organ is subject to many external cues [1, 62].

As in the brain, the genome of the SE displays a clear accumulation of non-CpG methylation on an organ-wide level. The similarity of the methylation patterns of the inner ear SE and brain may support the results of fate mapping experiments that suggest a dual origin of the inner ear, separate from that of the sensory placode endoderm and neuroepithelial cells [63]. This new paradigm suggests that not only does the otic placode originate from the ectoderm layer, and give rise to inner ear labyrinth cell types and the cochleovestibular ganglion, but that cells from the neural tube can also contribute directly to developing cells in the otic vesicle. An example of this crosstalk is the effect of Notch-signaling mitogens, which mark the prosensory side on the posterior-anterior axis and affects prosensory and proneuronal differentiation by modulating genetic regulatory networks [1, 64]. Newly characterized *Eya1*^*+*^ progenitor cells residing in the mammalian cochlea could give rise to spiral ganglion neurons (SGNs), as well as SE hair- and supporting cells, indicating a two-way direction, or plasticity, between the neuro- and non-neuronal portions of the inner ear and SGN [65], supported by the observation of nonCpG methylation accumulation in the adult inner ear SE. The interaction between the brain and the inner ear is further supported by the recent demonstration that auditory cortex interneuron development requires cadherins to operate in hair-cell mechanoelectrical transduction [66]. It is tempting to suggest that epigenetic regulation and fine tuning of gene expression is involved in directing the tissue’s hierarchical differentiation from the neuronal side of the inner ear to form the sensory organ.

Such information about how inner ear neuronal-sensory inter-tissue communication and cell-to-cell mitogen-induced epigenetic reconfiguration drive development and maturation of the organ, will be of great importance in the development of any treatment for hearing impairment. Significant therapeutic progress will require an improvement of *in vitro* differentiating mechanosensory progenitors that lack the natural 3D complexity and will necessitate an understanding of the need for a cell type-specific differentiation program, in combination with the correct environment for epigenetic reconfiguration. For *in vivo* work, epigenetic editing of various cellular populations involved in inner ear transcriptomic regulation might be crucial for establishing the conditions to achieve a true and efficient regeneration of the organ of Corti.

### From DNA methylation status to putative regulatory elements governing tissue development

Our analysis of the SE methylome reflects the DNA methylation status of a mixed population of cells. Nonetheless, our results offer an unprecedented opportunity obtain new insights into key pathways, biological and molecular processes crucial for SE development and maturation. We have used the single base resolution methylome to annotate putative regulatory elements, marked as UMRs or LMRs. By further analyzing their associated target gene-enriched GO terms and *in silico* predicted TF binding sites, we built an integrated regulatory network in the developing inner ear SE as a whole, or add new modules of regulation to known regulatory relationships between genes or pathways. Such relevant examples are the identification of deafness hubs within our *in silico* regulatory networks, focused around TFs such as *Sox2* and *Atoh1*; the identification of potentially novel regulators in the inner ear SE; and confirmation or expansion of poorly understood effectors of biological processes in the inner ear SE, such as FoxO1. Another example is the Bach2 TF, which is expressed in the inner ear SE and is an effector in oxidative stress-induced apoptosis [67, 68]. Taken together with the enrichment of the biological process ‘regulation of apoptotic process’ (GO:0042981) with various types of putative regulatory elements and potentially regulated genes indicated in our methylome, may hold potential for better prevention of apoptosis of inner ear SE cells and resulting hearing loss.

Within the networks for both DevTrans and MatTrans transitions, we could identify known deafness genes as upstream regulators likely bound to regulatory elements and as the downstream targets. In addition to providing a gene regulatory understanding of these essential hearing genes, the restriction of this analysis to DMRs illustrates the importance DNA methylation dynamics may play in the progression of inner ear SE development and hearing onset. This process is likely dependent on regulation of DNA methylation through DNA Methyltransferases (Dnmts), as knockdown of *Dnmt3a* was linked to impaired development of the inner ear in zebrafish^51^. Another key DNA methylation gene, *Dnmtl*, harbors a mutation linked to hereditary sensory and autonomic neuropathy type 1 (HSAN1E; OMIM 614116) [69] and autosomal dominant cerebellar ataxia, deafness, with narcolepsy (ADCA-DN; OMIM 604121), both involve deafness with late onset, suggesting a close connection between DNA methylation and the maintenance of tissue function and not necessarily the correct formation of the tissue.

Like the brain, the inner ear has been considered to be an immune privileged organ, lacking lymphatic drainage, and with the blood-labyrinth barrier, maintaining a separation between the circulation and the cochlea. This perspective changed when immune-reactive cellular components, such as resident tissue macrophages, were identified in the inner ear under non-inflammatory conditions [70, 71]. The methylome data we present point towards the presence and possible role of the immune system, or immune-related processes and signaling pathways, in the development and maturation of the inner ear SE. The association of immune-related processes to the sets of putative enhancers was especially compelling as it suggests a connection between inner ear immunity and either development or maturation, a possibility already proposed by ourselves and others. Inflammatory and immune responses may have a degenerative effect on several components of the inner ear [72-74]. We suggest that further study of the connection of annotated regulatory elements to immune-related genes, possibley NF-**k**B regulated genes in response to inflammation and noise damage, and the DNA methylation status of the elements, holds potential for developments in the field of regenerative medicine in the inner ear SE. Furthermore, we predict that cell type-specific methylome analysis of sensory versus non-sensory cells across the tonotopic map of the organ of Corti will reveal finite differential methylation that will enable us to explain the orchestration of topographic circuit assembly [75, 76], and may reveal positional epigenetic information carried by tissue embedded precursors that guide later differentiation processes.

### The mouse methylome is relevant for the human genome and deafness

Despite the increase in whole exome sequencing (WES) to determine the genetic cause of deafness, there are many individuals with hereditary hearing impairment whose cause of deafness remains unknown. Moreover, if WES does not determine the cause, the next step is to perform whole genome sequencing (WGS). The challenge is to determine if variants in noncoding regions are disease-causing, as there are only a few examples of such cases in the field of hearing impairment [77]. Epigenomic profiles, integrated with genetic data, may help define regulatory elements associated with organ development and gain of function, naturally marking them as loci with potentially pathogenic variants or altering activity of genes [49]. We have demonstrated the translation of regulatory data from the mouse SE model to human disease by taking mouse-to-human conserved putative enhancer elements (32-35% of mouse annotated regions), and cross-referencing them with validated deafness variants, GWAS and SNP datasets, available epigenetic data from numerous other tissues, and higher order 3D genome organization datasets. This analysis allowed us to identify SNPs and SNVs residing near putative enhancers as potential variants for hearing impairment. The example of Pendred syndrome and *SLC26A4* presented is one demonstration of this application. As a result, the inner ear SE methylome opens up a variety of possible targets for improved diagnostics, screening and study of various hearing impairments, otherwise unresolved.

We have presented a validation pipeline for one example of our methylome data, utilizing CRISPRon molecular technology, which can be applied on any mouse-to-human lifted over candidate enhancer sequence. In our exemplary candidate, chosen from the human GJB2- GJB6 locus, we have achieved several goals. First, we have demonstrated the chosen candidate enhancer regulates both GJB6 and the ncRNA (whose function is undefined) expression through what seems to be distal enhancer interactions. This interaction, positively modulating expression, is likely mediated by ATOH1, indicated by ATOHl-dependent activation of both GJB6 and the ncRNA. Moreover, an ATOH1 TFBS motif is present in the candidate enhancer sequence. Second, we have validated the value of the epigenetic exploratory phase of our work, performed in mice, and its translatability to human. Finally, we have presented the first CRISPRon-based modulation of endogenous inner ear relevant gene expression in cells that does not express them naturally.

## Conclusions

In this study, we have presented the first DNA methylome map in the mammalian inner ear SE. A genome-wide perspective of DNA methylation may guide inner ear lineage formation, tissue morphogenesis, and gain of auditory function. We used the methylation status and locus-specific methylation dynamics to annotate candidate genetic regulatory elements throughout the genome, presenting one of the first whole-genome regulatory maps for the developing hearing organ in mice. Our derived view on regulatory elements is reflective of the developing tissue as a whole, averaging the methylation status of all the cellular populations together. Future studies of DNA methylation in particular, and epigenetic regulation of the inner SE in general, should leverage this dataset to better understand specific pathway-related regulators or regulation of a specific cellular population compared to the entire organ of Corti. Regulatory elements studied in mouse models for deafness could potentially become crucial in the identification of non-coding variants causing NSHL, age-related hearing loss, and other types of hearing impairment in humans. In addition, they may serve as additional filtering criteria for the large numbers of SNVs resulting from whole-genome sequencing. Finally, a better understanding of the role of a global epigenetic regulatory process, as well as the dynamics of regulatory elements, may advance regenerative research of the inner ear sensory organ by directing the manipulation of gene expression, or repurposing currently available epigenetic therapeutics to treat or prevent the onset of a hearing impairment.

## Methods

### Animals and SE preparation

All procedures involving mice met the guidelines described in the National Institutes of Health Guide for the Care and Use of Laboratory Animals, and were approved by the Animal Care and Use Committees of Tel Aviv University (M-13- 114, M-13- 115 and 01-16- 100). C57BL/6J mice were purchased from Envigo, Jerusalem, Israel. The ages of the mice used were as follows: embryonic development day 16.5 (E16.5); post-natal, newborn on the day of birth (P0); post-natal, adult mice at 3 weeks old (P22) after hearing onset. A total of 6-8 mice were sacrificed by CO2 suffocation of the pregnant dam or the post-natal mice, followed by immediate decapitation (only decapitation was employed when dissecting newborns) for each replica. The skin at the top of the head was removed and the skull was opened at the dorsal portion along the midline. The brain tissue was removed and the auditory nerve was pulled out and cut. The inner ear, containing both the cochlea and the vestibule, was removed from the temporal bone, and placed in phosphate buffered saline (PBS). The cochlea was further dissected; the otic capsule, the spiral ligament and the stria vascularis were removed to expose the SE. Next, the SE was separated, base to apex, from the spongy modiluse bone.

### Whole genome bisulfite sequencing

We performed a genome wide analysis of the inner ear organ of Corti using an adaptation of the MethylC-Seq protocol [78]. gDNA was extracted from a pooled sample of 6-8 mice (12-16 sensory epithelium) using the QIAamp DNA Micro gDNA Kit (Qiagen#56304). Each sample was spiked with 0.5% of total gDNA mass, Lambda unmethylated DNA to serve as an internal control for bisulfite (BS) conversion rates. gDNA samples were fragmented using COVARIS S220 with Snap-Cap microTUBE with AFA fiber 6x16mm (Covaris #520045). Illumina-compatible NGS libraries were produced using the NEBNext^®^ Ultra DNA Library Prep Kit for Illumina^®^ (NEB#7370) and Multiplex Oligos for Illumina^®^ (Methylated Adaptor, Index Primers Set 1) (NEB#7535S). Adaptors were attached and the fragmented gDNA was cleaned and size selected using Agencourt AMPure XP (Beckman Coulter #A63881) magnetic beads. BS conversion was performed using Methylcode Bisulfite Conversion Kit (ABI #MECOV50). Each sample was spiked with 0.5% of total gDNA mass, Lambda unmethylated DNA to serve as an internal control for BS conversion rates. The initial prep was PCR amplified using KAPA HiFi Uracil+ polymerase with the index and universal Illumina compatible oligos, for 4-6 cycles. The library PCR was clean and size selected on AMPure XP beads. Library concentration was validated using the Qubit dsDNA High Sensitivity platform, and size distribution was tested using the DNA High Sensitivity Kit (Agilent#5067-4626). NGS was performed at BGI, China on an Illumina HiSeq 2500. Libraries were divided equally across lanes to minimize technical bias during sequencing.

### Methylation data analysis

WGBS reads were aligned to the mouse reference genome (mm10) using Bismark (version v0.14.0) [79] with default options. The bam files generated using Bismark were processed via samtools (0.1.18) [80] to remove duplicates. The library fold coverage was computed by dividing the number of uniquely aligned reads by the size of the genome. The mapping efficiency was computed by Bismark using the number of unique paired end alignments against the total paired end reads. The bisulfite conversion rates were computed by Bismark after mapping each sample with the lambda phage genome. The identification of the DMRs was performed using methylPipe (version 1.4.5) [81] only for CG context: 30% methylation difference, p-value < 0.05 non-parametric wilcoxon test, corrected for multiple testing. LMRs and UMRs were called as defined [18], using the R package MethylSeekR. Within the E16.5 and P22 time point combined libraries, we found that several genomic ranges contained an overlap between annotated UMRs and LMRs. We removed these ambiguous regions from our dataset.

### Gene ontology analysis

TSS proximal regulatory features were analyzed for GO terms and KEGG pathway using DAVID [82] (https://david.ncifcrf.gov/) with default parameters. Terms were ranked according to the P value (EASE score), which is derived from a modified Fisher’s exact test. Associated genes were determined according to previously described genomic annotation of the TSS proximal genomic ranges datasets, derived from lists of official gene symbols or ENTERZ gene id numbers (for list >3000 gene long).

For TSS distal elements, analysis of GO term enrichment was executed using the Homer suite (annotatePeaks.pl hg19 -go) [83] for DMRs or GREAT [84] (http://bejerano.stanford.edu/great) for putative regulatory elements characterized as tsLMRs. Terms were ranked according to the Binomial test Q score (Binom FDR Q-Val < 0.05) and the Hypergeometric test Q score (Hyper FDR Q-Val < 0.05).

### Transcription factor binding site (TFBS) motif prediction

Motif analysis was carried out using the Homer suite [83], which allowed us to analyze a large set of genomic ranges. The detected motifs were submitted for further processing only if the E-value <0.05. TF binding motifs were ranked according to their q-value; with the threshold was set to q-value < 0.05. TFBS were filtered in accordance with their expression levels in the inner SE RNA-seq available databases [6, 8].

### Identification of proxy SNPs and LMR liftover

SNPs for available hearing-related impairment were downloaded from the NHGRI-EGI GWAS Catalog (https://www.ebi.ac.uk/gwas/). LD SNPs were determined using rAggr, max distance 500kb (http://raggr.usc.edu). R-squared threshold of 0.5 was used based on previous studies showing enrichment at distal c/s-regulatory elements for SNPs from r2 1.0 to 0.5 [85, 86]. Mouse mm10 coordinate LMRs were converted to human hg19 coordinates using the UCSC LiftOver tool (https://genome.ucsc.edu/cgi-bin/hgLiftOver).

### CRISPRon system

We used the the pAC154-dual-dCas9VP160-sgExpression, a gift from Rudolf Jaenisch (Addgene plasmid # 48240) [53], and designed gRNAs using the CHOPCHOP web tool (http://chopchop.cbu.uib.no/) [87] to choose the top ranking guides. Oligos suitable for cloning were ordered from IDT according to cloning into the *Bbs*I site [53]. The guide sequences: gRNA#1 FWD 5’-caccG**acttcgcgatttttgcagag**-3’, gRNA#1 REV 5’-aaac**ctctgcaaaaatcgcgaagt**C-3’, gRNA#2 FWD 5’-caccG**gtaaaaacgtgagcgcgag**-3’, gRNA#2 REV 5’-aaac**ctcgcgctcacgtttttac**C-3’. Correct integration of guide RNAs into the plasmid was validated using Sanger sequencing with a sequencing primer from the U6 promoter upstream of the integration site, U6_SEQ 5’CAAGGCTGTTAGAGAGATAA-3’. All cloning design was done over the Benchling platform (www.benchling.com). Plasmids were harvested using NucleoSpin MidiPrep (MN#74010). Plasmids were transfected into 50% confluence cells seeded 18 hours before in 6-well plates. Transfection was done using JetPEI reagent according to the protocol (Polyplus#101). Cells were harvested for RNA 48 hours post transfection using the ZYMO Direct-zol RNA Miniprep Kit (Zymo research# R2070). RNA was measured using the NanoDrop and 500ng of RNA was taken from each sample to prepare cDNA using the qScript Reverse Transcription Kit (Quantabio 95047). All was performed in two technical repeats of each sample and across 5 biological replicates.

### Real Time qPCR

mRNA expression was evaluated using the PerfeCTa SYBR^®^ Green FastMix (QuantaBio #95074) in the StepOneTM Real-Time PCR System (Applied Biosystems). We designed primers using Primer3plus (http://primer3plus.com/cgi-bin/dev/primer3plus.cgi) with the default parameters to reach an amplicon of 80-150bp. Oligos were ordered from IDT in a 100uM stock. Primer sequences: GJB2_FWD 5'AAAAGCCAGTTTAACGCATTGC'3, GJB2_REV 5'TTGT GTT GGGAAAT GCTAGCG'3, GJB6_FWD 5'TGGCAAATTTGT GAACTGTCATG'3, GJB6_REV 5'TC AGTTGTTTGCAAT GATT GGC'3, RP11- 264J4.10_FWD 5'TGCTCAT GAAGAGGCAAAGC'3, RP11-264J4.10_REV 5'TTAACAAGCCGACTCAGCAC'3. All samples were normalized to GAPDH endogenous expression with the following primers GAPDH 5' GGAGCGAGATCCCTCCAAAAT'3 and GAPDH_REV 5'GGCTGTTGTCATACTTCTCATGG'3. The negative control was set as Non-Template Control (NTC). The relative expression level was measured using the 2-ddCt method. mRNA level from un-transfected cells (termed "noDNA") was set as 1. The data figure is presented as the mean±SE.

### Data retrieval

The data from this study have been deposited in NCBI Short Read Archive (SRA). The SRA accession number for the data generated from mouse inner ear SE is SRP111167, which will become public upon acceptance. Access to SRA metadata can be found at: ftp://ftp-trace.ncbi.nlm.nih.gov/sra/review/SRP111167_20170711_120631_679f47e946645e35bc0d606f07a1217e.

### Additional files

**Additional file 1**: Figure S1. Inner ear SE methylome characteristics. **a** The SE (right) is dissected from the cochlea of the inner ear (left). Scale bar: 50 μm. **b** The table shows number of mapped reads (in millions) and genomic coverage for the combined replicates at each time point (two replicates per time point). **c** Density plots showing the methylation level for UMRs and LMRs at each time point. E16.5, green; P0, orange; P22, blue. **d** Density plots showing width for UMRs and LMRs for each time point. **e-f** Pie charts showing the distribution of UMRs (**e**) and LMRs (**f**) across genomic features. **g-h** Bar charts showing the overlap of UMRs (**g**) and LMRs (**h**) with CpG islands (CGIs).

**Additional file 2**: Table S1. UMR and LMR coordinates.

**Additional file 3**: Table S2. UMR and LMR interacting genes.

**Additional file 4**: Table S3. UMR and LMR interacting deafness genes.

**Additional file 5**: Table S4. Homer TFBS analysis.

**Additional file 6**: Fig. S2. DMR general characteristics. **a** Violin plot presenting the distribution of % methylation difference (either hypo- or hypermethylated) at both transitions. Figure shows the median (internal black dot) and SE (internal black lines). **b** Density of DMR width (bp). **c** Genomic distribution of UMRs according to mm10 UCSC KnownGene database; DMR category (DevTrans/MatTrans Hypo-/Hypermethylated) is in line with (**b**), DMR overlapping to UMR (black), LMR, both, or none.

**Additional file 7**: Table S5. DMR coordinates.

**Additional file 8**: Fig. S3. Expanded MatTrans regulatory network. *In silico* transcriptional regulatory network based DMR target gene interactions during MatTrans. TFs with motifs enriched at DMRs are shown in yellow and TGs in blue.

**Additional file 9**: Table S6. DMR TF motifs and interacting genes.

**Additional file 10**: Table S7. GO analysis for regulatory networks.

**Additional file 11**: Fig. S4. Dynamic changes in UMRs and LMRs. Sankey plot visualization of a 3-way intersection between the complete subset of UMRs at the three time points examined presenting all UMRs (**a**) and LMRs (**b**). Dark grey connections between nodes indicate regions that either maintained or gained their UMR/LMR status during the transition. Light grey connections between nodes indicates regions that gain methylation through the transition and are no longer defined as UMRs (**a**), or lose or gain methylation and are no longer defined as LMRs at the end of the transition (**b**).

**Additional file 12**: Table S8. Time-specific UMR GO analysis.

**Additional file 13**: Table S9. Time-specific LMR GO analysis.

**Additional file 14**: Fig. S5. Analyzing the correlation between the DNA methylation dynamics and transcriptome of the inner ear SE based on the SHIELD database [6]. Scatter plot of methylation difference [%] against gene expression [Log2(Fold Change)] of promoters from DevTrans (**a**) and MatTrans (**b**) combined hair cells and surrounding cell gene expression. Scatter plot of methylation difference [%] against gene expression [Log2(Fold Change)] of genebody from DevTrans (**c**) and MatTrans (**d**) combined hair cells and surrounding cell gene expression.

**Additional file 15**: Table S10. GO analysis of genes presenting expected correlation between DNA methylation and gene expression.

## Abbreviations

SE: sensory epithelium
E: embryonic day
P: postnatal day
ES: embryonic stem
UMR: unmethylated regions
LMR: low-methylated regions
DHS: DNase I hypersensitive sites (DHS)
TSS: transcription start sites
CGI: CpG island
TF: transcription factor
TFBS: transcription factor binding sites
3D: three-dimensional
DevTrans: developmental transition
MatTrans: maturation transition
DMR: differentially methylated region
TG: target gene
GO: Gene Ontology
LD: linkage disequilibrium
EVA: enlarged vestibular aqueduct
NHEK: Normal Human Epidermal Keratinocytes
gRNA: guide RNAs
WES: whole exome sequencing
WGS: whole genome sequencing

## Declarations

### Ethics approval

All procedures involving mice met the guidelines described in the National Institutes of Health Guide for the Care and Use of Laboratory Animals, and were approved by the Animal Care and Use Committees of Tel Aviv University (M-13- 114, M-13- 115 and 01-16- 100).

### Consent for publication

Not applicable

### Availability of data and materials

The datasets supporting the conclusions of this article are available in the NCBI Short Read Archive (SRA). The SRA accession number for the data generated from mouse inner ear SE is SRP111167, which will become public upon acceptance. Access to SRA metadata can be found at: ftp://ftp-trace.ncbi.nlm.nih.gov/sra/review/SRP111167_20170711_120631_679f47e946645e35bc0d606f07a1217e, and within the article and its additional files.

## Author contributions

O.Y.-B., K.B.A., and R.D.H. conceived the project and together with C.V., designed experiments, analyzed the data, and wrote the manuscript, with input from all authors. K.K., N.D.J., C.A., M.P., O.Y.-B., C.V., and K.P. performed the bioinformatics and data analysis.

O.Y.-B, C.V., T.K.-B., K.U., Y.N., and Y.B. performed the laboratory experiments. All authors have read and approved the manuscript for publication.

## Acknowledgements

This work was supported by the United States-Israel Binational Science Foundation grant no. 2013027 (K.B.A. and R.D.H.); the Israel Science Foundation grant no. 2033/16 (K.B.A.); the National Institutes of Health/NIDCD R01DC011835 (K.B.A.); NIH/NIAMS R01AR065952 (R.D.H.); NIH/NIDDK R01DK103667 (R.D.H.); Action on Hearing Loss Flexi Grant (K.BA.) and I-CORE Program of the Planning and Budgeting Committee and The Israel Science Foundation no. 41/11 (K.B.A.). K.B.A. is an incumbent of the Drs. Sarah and Felix Dumont Chair for Research of Hearing Disorders. This work was performed in partial fulfillment of the requirements for a Ph.D. degree by Ofer Yizhar-Barnea, recipient of the Prof. Dr. Heinrich Neumann von Hethars Doctoral Scholarship, at the Sackler Faculty of Medicine, Tel Aviv University, Israel.

